# Generation of knock-in Cre and FlpO mouse lines for precise targeting of striatal projection neurons and dopaminergic neurons

**DOI:** 10.1101/2025.06.15.659794

**Authors:** Eddy Albarran, Akira Fushiki, Anders Nelson, David Ng, Corryn Chaimowitz, Laudan Nikoobakht, Tanya Sippy, Darcy S. Peterka, Rui M. Costa

## Abstract

The basal ganglia and midbrain dopaminergic systems are critical for motor control, reward processing, and reinforcement learning, with dysfunction in these systems implicated in numerous neurodegenerative and neuropsychiatric disorders. To enable precise genetic targeting of key neuronal populations, we generated and characterized five knock-in mouse lines: *Drd1-Cre, Adora2a-Cre, Drd1-FlpO, Adora2a-FlpO*, and *DAT-FlpO*. These lines allow for Cre-or FlpO-mediated recombination in dopamine D1 receptor-expressing spiny projection neurons (SPNs), adenosine A2a receptor-expressing SPNs, and dopamine transporter (DAT)-expressing neurons in the midbrain. Histological analyses confirmed recombinase activity in expected brain regions, and whole-cell electrophysiological recordings validated the intrinsic excitability profiles of each neuronal subpopulation. These tools provide high specificity and reliability for studying basal ganglia circuitry and dopaminergic neurons. By enabling targeted manipulations, these openly available knock-in lines will advance research into the neural mechanisms underlying motor control, reward, and neuropsychiatric diseases.

## INTRODUCTION

The basal ganglia are a collection of interconnected subcortical nuclei that play critical roles in motor control, reward processing, and decision-making (Grillner and Robertson, 2016; Yin, 2017; Arber and Costa, 2022). Dysfunctions within the basal ganglia have been implicated in a range of neurological and psychiatric disorders, including Parkinson’s disease, Huntington’s disease, and addiction (Foerde and Shohamy, 2011; Rice et al., 2011; Calabresi et al., 2014; Gunaydin and Kreitzer, 2016). Central to basal ganglia function are two major neuronal populations: the dopamine D1 receptor-expressing SPNs and the dopamine D2 receptor-expressing SPNs, canonically postulated to give rise to the direct and indirect pathways of the basal ganglia. These populations are modulated by input from dopamine transporter (DAT)-expressing midbrain neurons located in the ventral tegmental area (VTA) and substantia nigra pars compacta (SNc), which provide critical modulatory input to the basal ganglia. Together, these neuronal populations form the foundation for many of the basal ganglia’s key functions and their dysfunction in disease.

Genetically engineered mouse models have transformed neuroscience research, enabling cell type-specific manipulation of gene expression and activity. Among these tools, Cre and Flp recombinases have emerged as powerful drivers of conditional gene expression and deletion (Heintz, 2004; He et al., 2016; Pettibone et al., 2019; Kramer et al., 2021; Oppman et al., 2025). While traditional transgenic approaches utilizing bacterial artificial chromosome (BAC) constructs have been widely employed and have been very useful, expression does not always align with the endogenous expression of the genes in hand, and they can exhibit transgenerational discrepancies in expression (He et al., 2016). Knock-in strategies provide the ability to access cellular populations during development (Gerits et al., 2007) and provide superior specificity by aligning recombinase expression with endogenous gene regulatory elements (He et al., 2016). Such precision is especially critical for studying complex systems like the basal ganglia, where small variations in gene expression can lead to significant functional differences (Berke et al., 1998; Heintz et al., 2004).

Here, we describe the generation and characterization of five novel knock-in mouse lines: Drd1-Cre (D1-Cre), Adora2a-Cre (A2a-Cre), Drd1-FlpO (D1-FlpO), Adora2a-FlpO (A2a-FlpO), and DAT-FlpO. These lines were engineered to enable precise Cre-or FlpO-mediated recombination in D1R-expressing SPNs, A2aR-expressing SPNs, and DAT-expressing dopaminergic neurons. To validate the specificity and targeting of these lines, we performed histological analyses using reporter mice demonstrating recombinase activity in the corresponding neurons of the striatum, and in midbrain dopaminergic neurons. Additionally, electrophysiological recordings from D1R- and A2aR-expressing SPNs in the striatum and DAT neurons in the VTA and SNc confirmed the intrinsic excitability profiles expected from those cell populations.

By combining the specificity of knock-in recombinase targeting with rigorous expression and electrophysiological validation, our study validates these valuable targeted mouse lines for the neuroscience community. These lines will facilitate the investigation of basal ganglia circuitry and its role in both normal and pathological conditions and permit combinatorial targeting of specific populations. We hope that these widely available lines will facilitate our understanding of dopamine-driven reinforcement learning, motor control, and neuropsychiatric disease mechanisms.

## RESULTS

### Genetic strategy and generation of knock-in lines

Knock-in mouse lines were designed such that transcription of inserted recombinases would be controlled by endogenous promoter activity. A CRISPR-Cas9 approach was used to insert donor sequences containing T2A (Kanca et al., 2019) followed by Cre or FlpO recombinase sequences just before the stop codon of the endogenous gene, preserving gene function (Figure 1).

**Figure 1.**
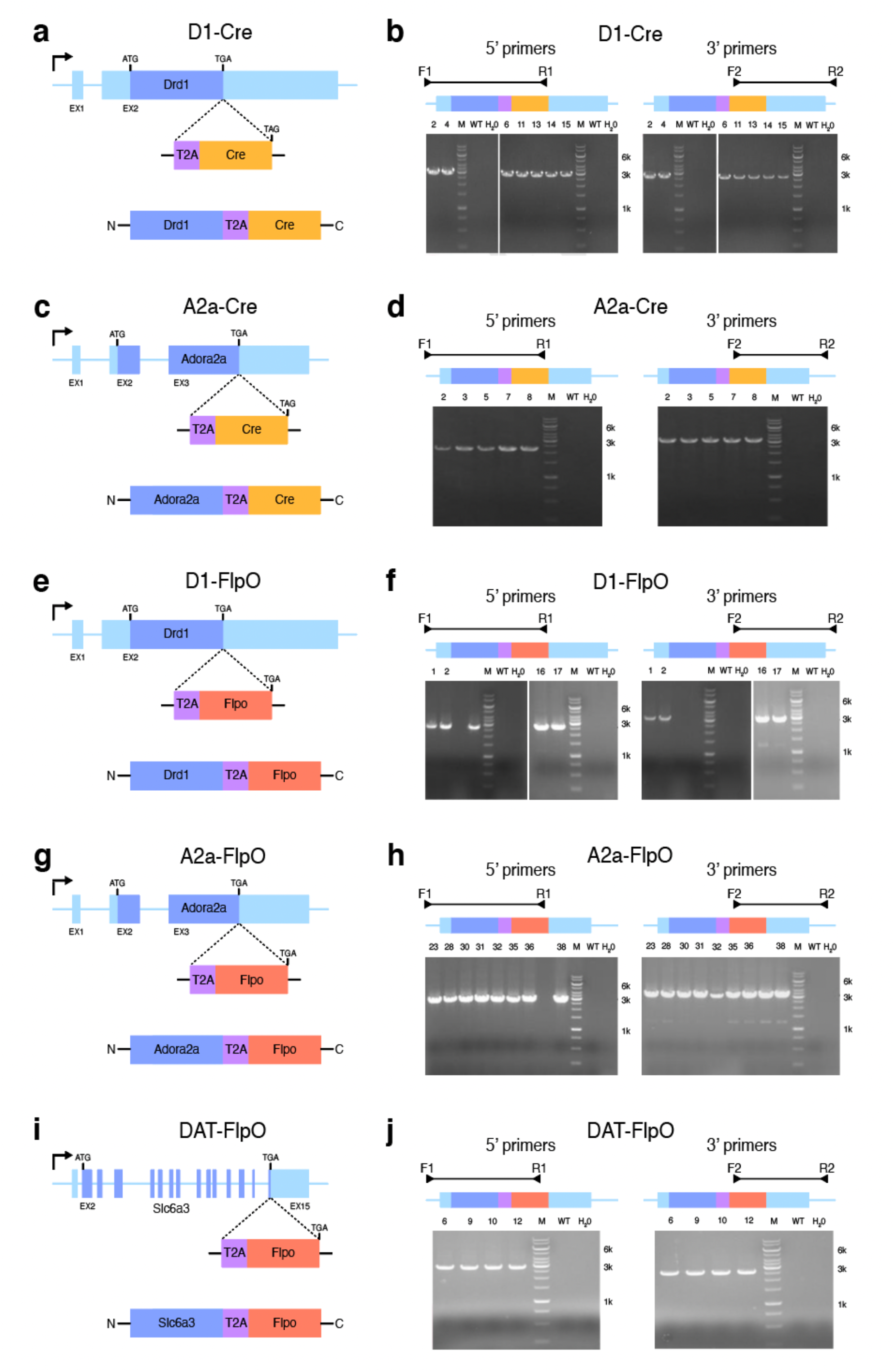
Genetic strategy for CRISPR-Cas9-mediated generation of knock-in lines. **a**, Schematic of generation of D1-Cre mice. (Top) *Drd1* gene with (middle) targeting vector replacing the stop codon with a T2A sequence and Cre recombinase resulting in (bottom) a modified locus expressing Cre under the control of the endogenous *Drd1* promoter. **b**, PCR-based genotyping strategy for D1-Cre mice using 5’ and 3’ primer sets. Positive bands denote presence of T2A-Cre insert in F1 mice, which are not present in wildtype (WT) or negative (H_2_O) controls. **c, d**, Genetic schematic (**c**) and PCR-genotyping strategy (**d**) for the generation of A2a-Cre mice, with insertion of a T2A-Cre sequence replacing the *Adora2a* stop codon. **e, f**, Genetic schematic (**e**) and PCR-genotyping strategy (**f**) for the generation of D1-FlpO mice, with insertion of a T2A-FlpO sequence replacing the *Drd1* stop codon. **g, h**, Genetic schematic (**g**) and PCR-genotyping strategy (**h**) for the generation of A2a-FlpO mice, with insertion of a T2A-FlpO cassette replacing the *Adora2a* stop codon. **i, j**, Genetic schematic (**i**) and PCR-genotyping strategy (**j**) for the generation of DAT-FlpO mice, with insertion of a T2A-FlpO sequence replacing the *Slc6a3* stop codon. M: DNA mass ladder.

A gRNA recognizing the mouse endogenous gene, donor DNA containing the “T2A-recombinase” cassette, and Cas9 mRNA were co-injected into fertilized C57Bl/6J mouse embryos to generate targeted CRISPR knock-in offspring. F0 founder animals were identified by PCR using insertion screening primers (Table 1) and confirmed by sequence analysis. Founders were then crossed to C57Bl/6J mice to obtain F1 pups and germline transmission of knock-in sequences were once again confirmed by PCR (Figure 1). The absence of random insertion was confirmed by vector backbone PCR screening.

**Table 1.**
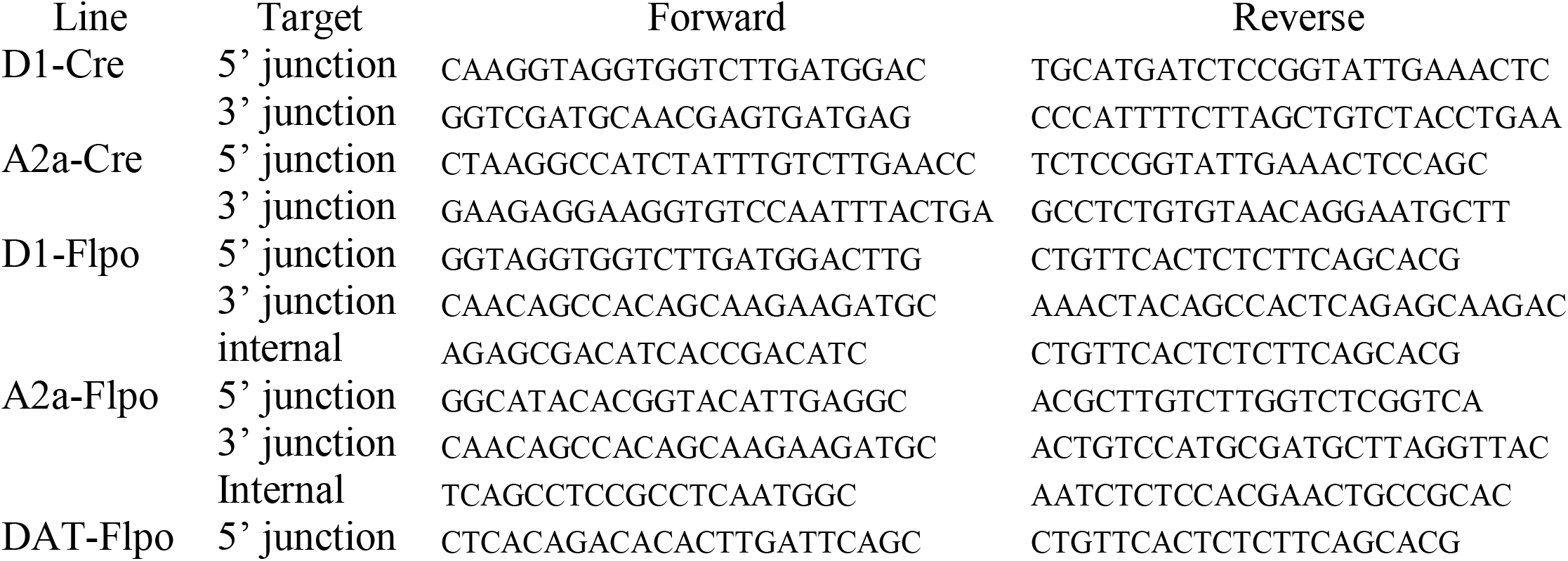
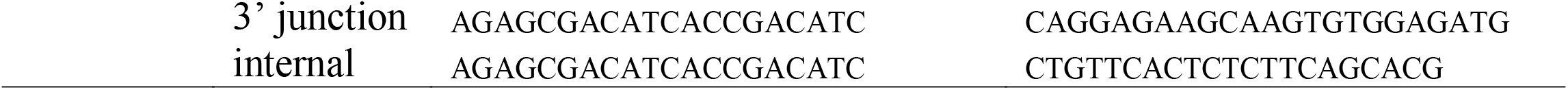
Screening primers.

### Whole-brain histology of D1 and A2a knock-in lines confirms specificity

A major motivation for generating knock-in mouse lines using the endogenous D1 and A2a receptor promoters was to gain specific genetic and anatomical access to the canonical direct and indirect pathways of the basal ganglia, respectively. To assess pathway expression specificity of these lines, we crossed the new D1-Cre and A2a-Cre lines with the Ai9 reporter line (Madisen et al., 2010), and crossed the D1-FlpO and A2a-FlpO lines to the RCE:FRT reporter line (Sousa et al., 2009), allowing for visualization of Cre recombinase activity through tdTomato expression and FlpO recombinase activity through EGFP expression.

D1-Cre;Ai9, D1-FlpO;RCE:FRT, A2a-Cre;Ai9, and A2a-FlpO;RCE:FRT mice were then perfused, and the brains removed and processed for immunofluorescence microscopy (Figure 2). Through examination of the fluorescent reporter signal in whole-brain sections, we confirmed that both the D1-Cre and D1-FlpO knock-in lines exhibited clear recombinase activity throughout the canonical direct pathway, including the striatum (Figures 2a-d,f-i), the globus pallidus internus (GPi) (Figure 2a,d,f,i), and the substantia nigra pars reticulata (SNr) (Figure 2a,e,f,j), but not in the globus pallidus externus (GPe) (Figure 2a,d,f,i). By contrast, the A2a-Cre and A2a-FlpO knock-in lines exhibited strong recombinase activity in the canonical indirect pathway, including the striatum (Figure 2k-n,p-s) and the GPe (Figure 2k,n,p,s), but not the GPI (Figure 2k,n,p,s) or the SNr (Figure 2k,o,p,t). Additionally, we observed more restricted reporter expression in the FlpO lines compared to the Cre lines, although this may be attributed to differences in reporter lines used. In combination, our histological experiments support that indeed the D1 and A2a knock-in lines exhibit expression specificity to the direct and indirect pathways as desired. Outside of the basal ganglia, we also noted lower Cre- and FlpO-dependent reporter expression in the cortex, thalamus, and hippocampus in the knock-in lines, which was to be expected (Levey et al., 1993; Heintz, 2004; Rani and Kanungo, 2006; Gangarossa et al., 2012).

**Figure 2.**
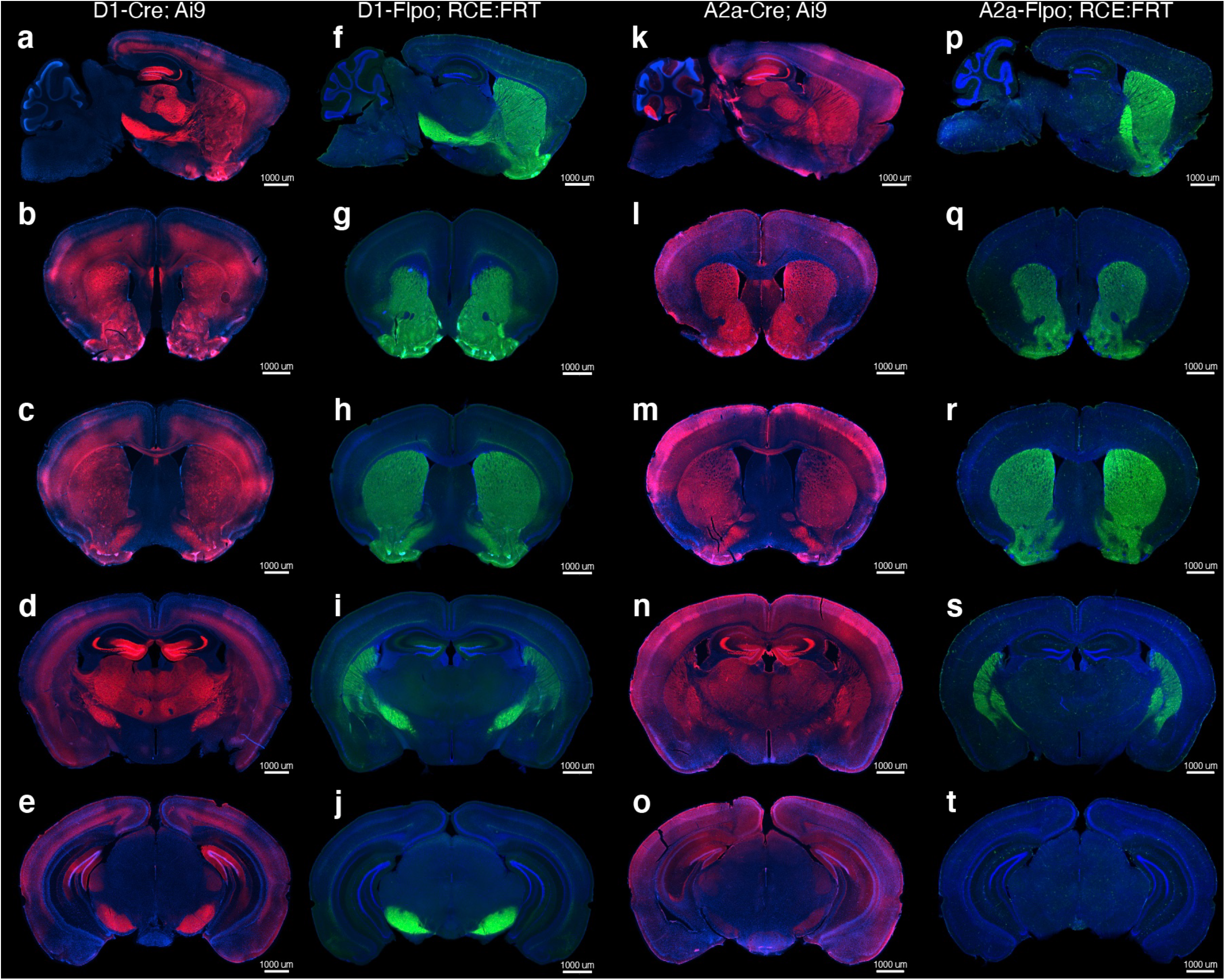
Whole-brain histological validation of D1 and D2 knock-in lines. **a-e**, Representative images of mouse brain sections from D1-Cre;Ai9 mice showing Cre-dependent tdTomato expression. **f-j**, Representative images of mouse brain sections from D1-FlpO;RCE:FRT mice showing FlpO-dependent EGFP expression. **k-o**, Representative images of mouse brain sections from A2a-Cre;Ai9 mice showing Cre-dependent tdTomato expression. **p-t**, Representative images of mouse brain sections from A2a-FlpO;RCE:FRT mice showing FlpO-dependent EGFP expression. Sagittal sections (**a, f, k, p**): ML +1.90. Coronal sections (**b, g, l, q**): AP +1.40. Coronal sections (**c, h, m, r**): AP +0.60. Coronal sections (**d, i, n, s**): AP -1.60. Coronal sections (**e, j, o, t**): AP -3.20. Scale-bar: 1000µm.

### DAT-FlpO knock-in line recombinase expression labels midbrain dopaminergic neurons

To confirm the restriction of FlpO recombinase to dopaminergic neurons in the new DAT-FlpO line, we crossed mice to the reporter line Ai65F (Daigle et al., 2018). We then perfused DAT-FlpO;Ai65F mice and performed immunohistochemical imaging of tdTomato and tyrosine hydroxylase (TH) to compare the whole-brain expression of FlpO recombinase and dopaminergic neurons respectively. We observed clear tdTomato expression in midbrain regions corresponding to VTA and SNc (Figure 3a), further corroborated by the strong overlap in TH immunoreactivity that is a hallmark of these dopaminergic structures (Figure 3a) (Daubner et al., 2011).

**Figure 3.**
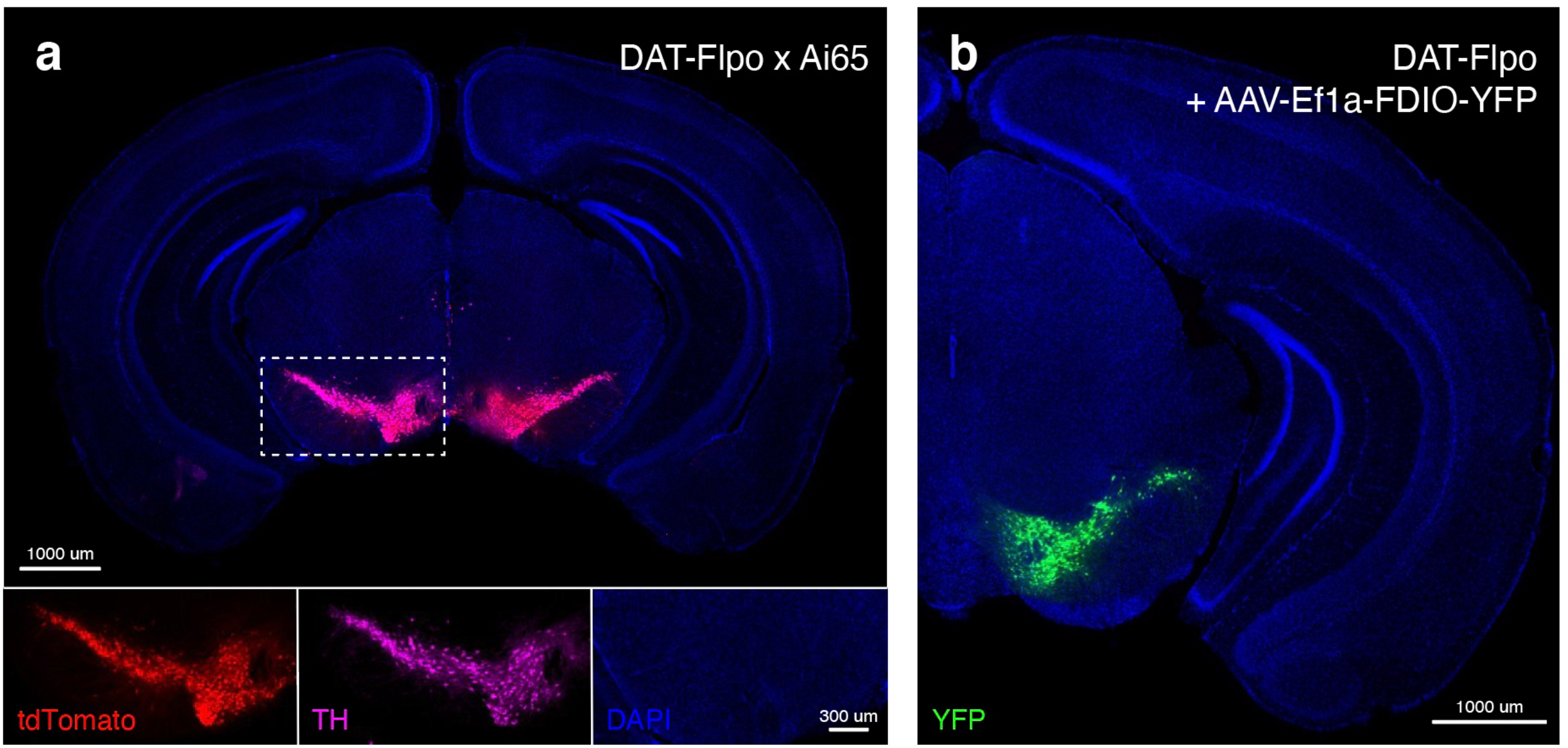
Histological validation of DAT-FlpO knock-in line. **a**, Representative coronal section from a DAT-FlpO;Ai65 mouse. TH antibody immunoreactivity (labeling midbrain dopamine neurons) significantly overlapped with FlpO-dependent tdTomato expression driven by the knock-in DAT-FlpO line. **b**, Viral expression of a FlpO-dependent YFP construct into the midbrain of DAT-FlpO mice confirms the restricted expression of YFP to midbrain VTA and SNc regions.

To further validate the utility of the DAT-FlpO line to target dopaminergic populations, we used a viral expression approach. We retro-orbitally injected DAT-FlpO mice with an adeno-associated virus (AAV) encoding a FlpO-dependent YFP construct. Four weeks post injection, we perfused these mice and observed clear YFP expression in VTA and SNc neurons, but not in the surrounding structures (Figure 3b). Combined with our histological experiments using reporter line crosses, our anatomical results using AAVs confirm that DAT-FlpO recombinase activity is clearly, and specifically, expressed in dopaminergic neurons.

### Electrophysiological characterization of D1 and A2a knock-in lines

Our histological experiments appeared to confirm specific recombinase activity in the desired neuronal populations, but to better assess if the knock-in lines targeted the distinct direct- and indirect-pathway SPN subtypes in the basal ganglia, we performed whole-cell patch clamp recordings of striatal SPNs in our D1 and A2a knock-in mouse lines. D1-Cre and A2a-Cre lines were crossed to the Ai9 reporter line and acute coronal brain sections prepared for acute *ex vivo* recordings of striatal SPNs. SPNs with visually-identified tdTomato-positive fluorescence in D1-Cre and A2a-Cre sections were classified as D1 and D2 receptor expressing SPNs, respectively (Figure 4a,b), with tdTomato-negative SPNs classified reciprocally as putative D2 and D1 receptor expressing SPNs, respectively. Increasing steps of current injections were used to quantify intrinsic electrical properties such as input-firing curves (Figure 4c-e) and I-V curves (Figure 4f). Comparing all recordings across D1-Cre and A2a-Cre lines, we observed no significant difference in resting membrane potential across groups (Figure 4g). As expected, we observed a clear difference in rheobase currents across groups, with putative D1-receptor expressing SPNs (D1-Cre + tdTomato-positive and A2a-Cre + tdTomato-negative) exhibiting significantly greater firing thresholds (310.48 ± 10.83 and 258.00 ± 13.37 pA) compared to putative D2-receptor expressing SPNs (A2a-Cre + tdTomato-positive and D1-Cre + tdTomato-negative) (176.25 ± 9.70 and 172.38 ± 10.37 pA) (Figure 4e,h). These results are consistent with observations of increased input resistance in indirect-pathway SPNs compared to direct-pathway SPNs (Al-muhtasib et al., 2018).

**Figure 4.**
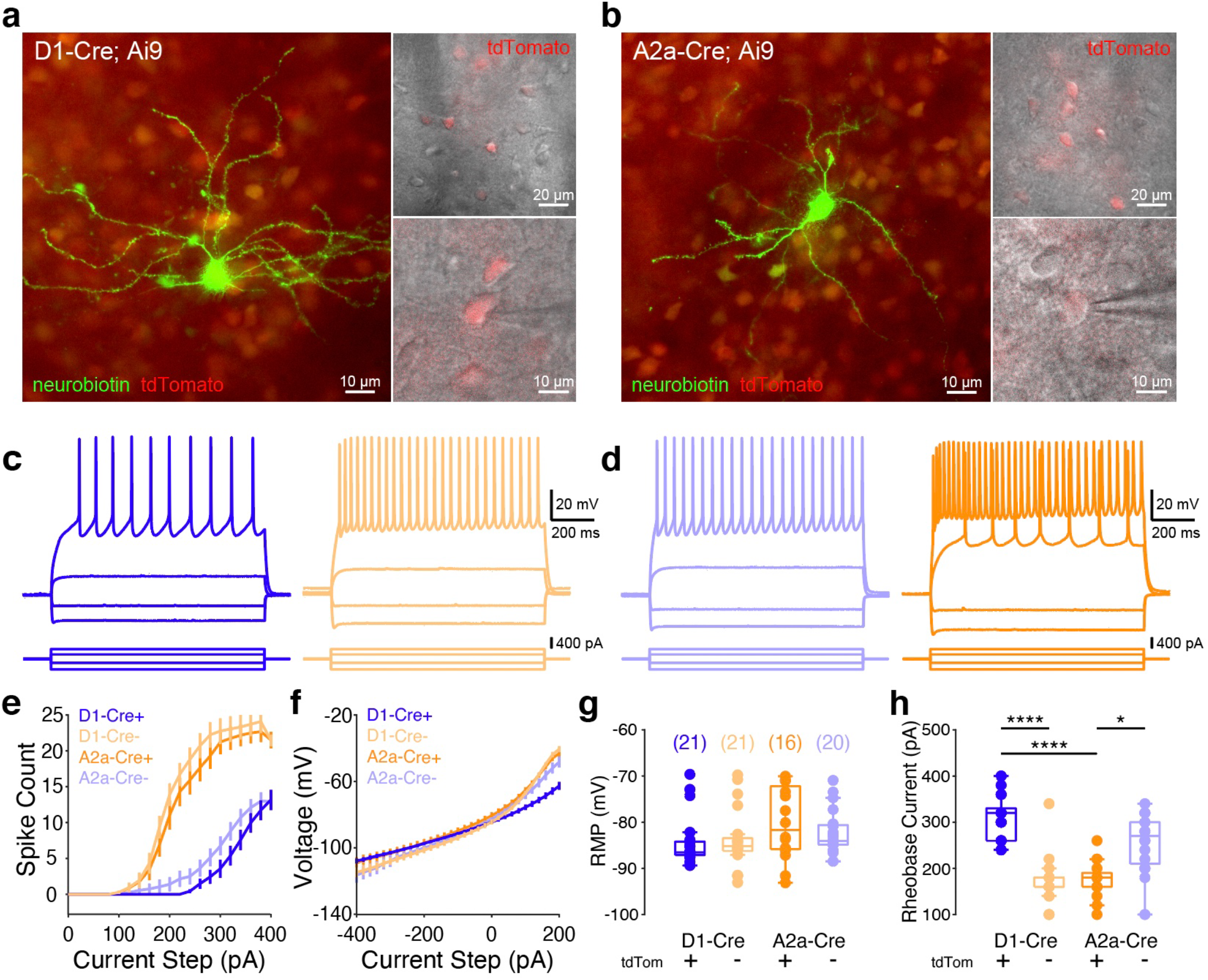
Electrophysiological validation of D1-Cre and A2a-Cre knock-in lines. **a**,**b**, Left: representative histological staining of neurobiotin-filled SPNs from whole-cell recordings from D1-Cre;Ai9 mice (**a**) and A2a-Cre;Ai9 mice (**b**). Right: DIC images of striatal SPNs, with tdTomato expression driven in D1 SPNs (**a**) or D2 SPNs (**b**). **c**,**d**, Representative whole-cell recording traces of SPNs with current injections evoking voltage responses and action potential firing. **e**, f-I curves quantifying SPN firing in response to injected current values. **f**, I-V curves quantifying SPN voltage changes in response to injected current values. **g**, Resting membrane potentials across all recorded SPNs (D1-Cre tdTom+ -84.34 ± 1.17 mV, n = 21 cells / 5 mice; D1-Cre tdTom--83.95 ± 1.40 mV, n = 21 cells / 5 mice; A2a-Cre tdTom+ -80.28 ± 1.94 mV, n = 16 cells / 5 mice; A2a-Cre tdTom--82.24 ± 1.05 mV, n = 20 cells / 5 mice). **h**, Rheobase currents for all recorded SPNs across D1-Cre and A2a-Cre animals (D1-Cre tdTom+ 310.48 ± 10.83 pA; D1-Cre tdTom-172.38 ± 10.37 pA; A2a-Cre tdTom+ 176.25 ± 9.70 pA; A2a-Cre tdTom-258.00 ± 13.37 pA). Significantly decreased rheobase currents in putative D2 SPNs (D1-Cre tdTom+ vs tdTom-: p < 0.0001; A2a-Cre tdTom+ vs tdTom-: p = 0.0142).

Next, we conducted whole-cell patch clamp recordings in D1-FlpO and A2a-FlpO knock-in lines. We crossed mice to the Ai65F reporter line and prepared acute coronal sections containing the striatum and recorded from tdTomato-positive and tdTomato-negative SPNs in both lines (Figure 5a-f). As with our Cre knock-in recordings, we observed no significant difference in resting membrane potentials across all recorded FlpO knock-in groups (Figure 5g). Similar to our findings in the Cre-lines above, the putative D1 receptor expressing SPNs (D1-FlpO + tdTomato-positive and A2a-FlpO + tdTomato-negative) once again clearly exhibited increased rheobase currents compared to putative D2 receptor expressing SPNs (A2a-FlpO + tdTomato-positive and D1-FlpO + tdTomato-negative) (Figure 5h). Our combined Cre and FlpO electrophysiological data suggest that the newly generated D1-Cre, A2a-Cre, D1-FlpO, and A2a-FlpO lines are specifically expressing recombinases in the direct (D1R-expressing) and indirect (D2R-expressing) pathways, consistent with our anatomical immunohistochemical data (Figure 2). Furthermore, our recordings validate that genetic knock-in of recombinase sequences did not fundamentally disrupt SPN electrophysiological properties.

**Figure 5.**
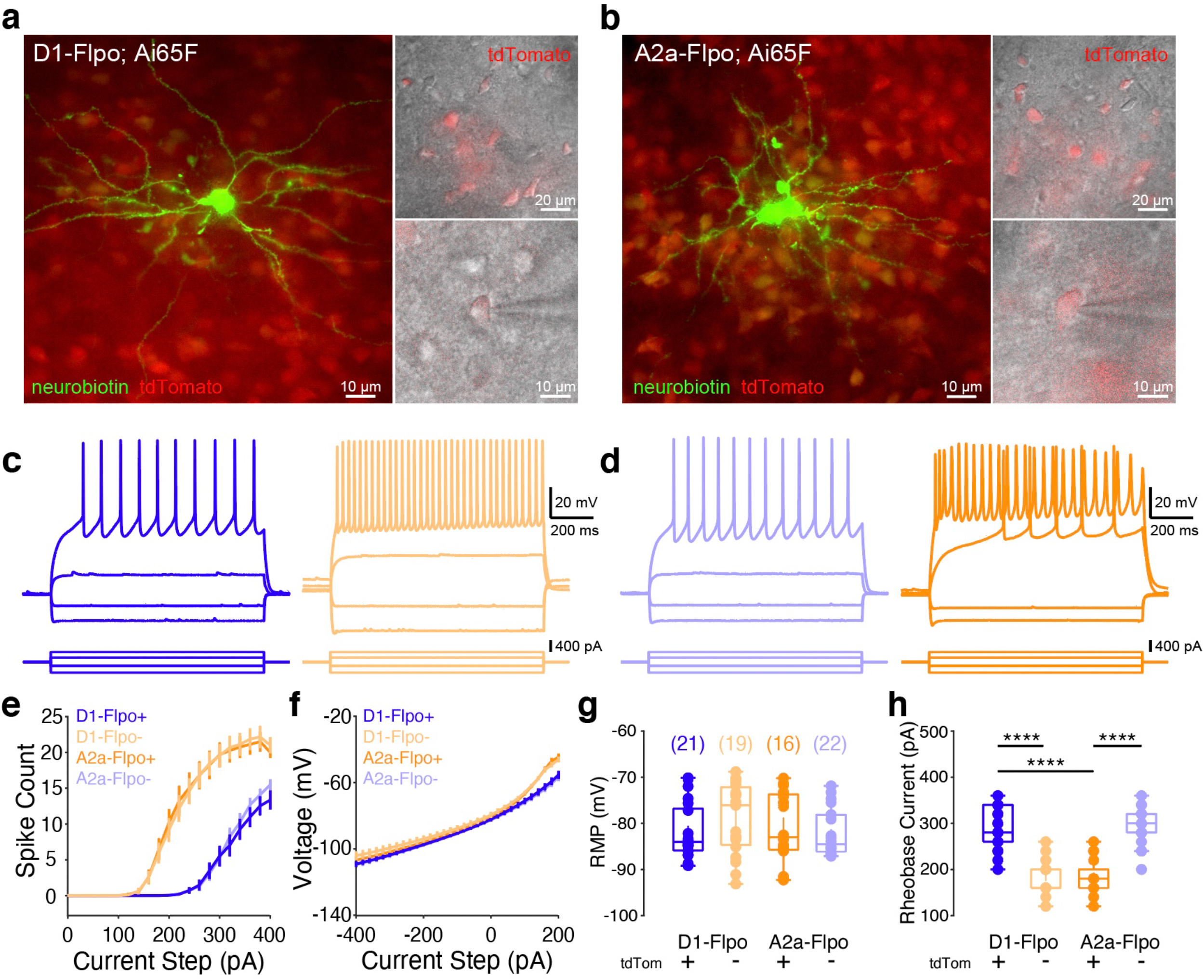
Electrophysiological validation of D1-FlpO and A2a-FlpO knock-in lines. **a**,**b**, Left: representative histological staining of neurobiotin-filled SPNs from whole-cell recordings from D1-FlpO;Ai65F mice (**a**) and A2a-FlpO;Ai65F mice (**b**). Right: DIC images of striatal SPNs, with tdTomato expression driven in D1 SPNs (**a**) or D2 SPNs (**b**). **c**,**d**, Representative whole-cell recording traces of SPNs with current injections evoking voltage responses and action potential firing. **e**, f-I curves quantifying SPN firing in response to injected current values. **f**, I-V curves quantifying SPN voltage changes in response to injected current values. **g**, Resting membrane potentials across all recorded SPNs (D1-FlpO tdTom+ -81.87 ± 1.30 mV, n = 21 cells / 5 mice; D1-FlpO tdTom--79.01 ± 1.77 mV, n = 19 cells / 5 mice; A2a-FlpO tdTom+ -80.60 ± 1.82 mV, n = 16 cells / 5 mice; A2a-FlpO tdTom--82.32 ± 1.05 mV, n = 22 cells / 5 mice). **h**, Rheobase currents for all recorded SPNs across D1-FlpO and A2a-FlpO animals (D1-FlpO tdTom+ 291.43 ± 9.72 pA; D1-FlpO tdTom-183.16 ± 8.27 pA; A2a-FlpO tdTom+ 183.75 ± 9.53 pA; A2a-FlpO tdTom-294.55 ± 7.92 pA). Significantly decreased rheobase currents in putative D2 SPNs (D1-FlpO tdTom+ vs tdTom-: p < 0.0001; A2a-FlpO tdTom+ vs tdTom-: p < 0.0001).

### Electrophysiological characterization of the DAT-FlpO knock-in line

Dopaminergic neurons of the midbrain are a particularly vulnerable population, with disruptions to their activity resulting in cell death (Hirsch et al., 1988; Damier et al., 1999). Because of this sensitivity, we aimed to confirm that dopaminergic neurons exhibited normal electrophysiological properties in the newly generated DAT-FlpO line. Crossing the DAT-FlpO line to the Ai65F reporter line (DAT-FlpO;Ai65F) allowed for labeling tdTomato-positive midbrain dopaminergic neurons for whole-cell patch clamp recordings from anatomically defined VTA and SNc (Figure 6a). Current step injections in recordings in VTA and SNc confirmed that tdTomato-positive putative dopaminergic neurons exhibit normal input-firing and I-V curves (Figure 6b-d), with no observed differences between VTA and SNc neurons in recording input resistance, resting membrane potential, or rheobase current (Figure 6e-h) (Margolis et al., 2006). These recordings validate that midbrain dopamine neurons in the DAT-FlpO knock-in line possess largely intact electrophysiological properties, with expected characteristics across all measurements of intrinsic excitability.

**Figure 6.**
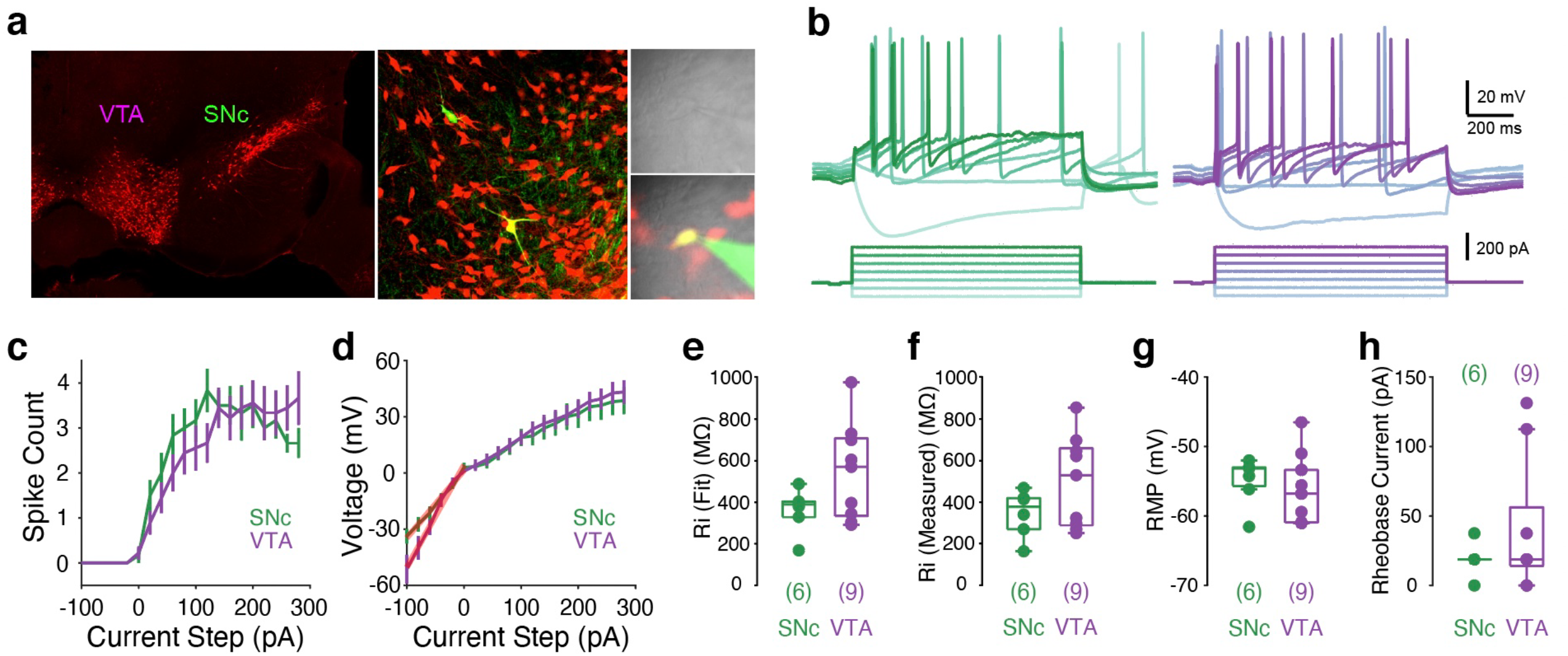
Electrophysiological validation of DAT-FlpO knock-in line. **a**, Left: histological section of DAT-FlpO;Ai65F brain containing VTA and SNc. Middle: post-recording histology of dopamine neuron filled with neurobiotin during whole-cell records. Right: DIC image (top) and epifluorescence image of tdTomato dopamine neurons (bottom) targeted for recordings (pipette solution containing alexa dye). **b**, Representative voltage traces of VTA (left) and SNc (right) neurons in response to current injections. **c**, f-I curves quantifying dopamine neuron firing in response to injected current values. **d**, I-V curves quantifying dopamine neuron voltage changes in response to injected current values (red line indicated linear fit). **e**, Input resistances across recorded dopamine neurons, based on linear fits of I-V curves (SNc: 360.37 ± 43.76 MΩ, n = 6 cells; VTA: 544.97 ± 77.39 MΩ, n = 9 cells). **f**, Directly measured input resistances (SNc: 344.90 ± 46.11 MΩ; VTA: 497.60 ± 73.32 MΩ). **g**, Measured resting membrane potentials across all dopamine neurons (SNc: -55.03 ± 1.43 mV; VTA: -56.18 ± 1.70 mV). **h**, Measured rheobase currents across all dopamine neurons (SNc: 20.00 ± 5.16 pA; VTA: 42.22 ± 17.14 pA).

## DISCUSSION

In this study, we generated and characterized five novel knock-in mouse lines – D1-Cre, A2a-Cre, D1-FlpO, A2a-FlpO, and DAT-FlpO – that enable precise genetic targeting of key neuronal populations in the basal ganglia and midbrain dopaminergic system. By leveraging CRISPR-Cas9-mediated knock-in strategies, we ensured that Cre and FlpO recombinases were expressed under the control of endogenous promoters, thereby preserving the fidelity of gene expression patterns, allowing for cell type-specific genetic manipulations. Our histological and electrophysiological analyses confirmed that these lines exhibit high anatomical and functional specificity, offering valuable tools for studying basal ganglia and dopamine neuron circuitry and function.

Our histological validation demonstrated that recombinase expression closely matches expected anatomical distributions of D1R-, A2aR-, and DAT-expressing neurons, confirming the accuracy of our knock-in strategy. Importantly, while recombinase expression was also observed outside the basal ganglia (e.g., cortex, thalamus, hippocampus), these findings are consistent with known endogenous expression patterns of D1R, A2aR, and DAT (Levey et al., 1993; Lammel et al., 2015). Although it is known that genetic modifications – particularly those involving recombinase integration – can disrupt neuronal physiology (Bäckman et al., 2006; O’Neill et al., 2017), our electrophysiological analyses of striatal SPNs and midbrain dopaminergic neurons confirmed that recombinase expression does not significantly alter intrinsic excitability properties, supporting the functional integrity of the knock-ins.

Future studies will be able to utilize these generated knock-in lines to gain genetic access to basal ganglia and dopamine neurons with greater precision. Indeed, combining these Cre and FlpO lines with the growing toolkit of Cre- and FlpO-dependent reporter lines, optogenetic and chemogenetic constructs, and optical sensors, will allow for greater investigation of circuitry, function, and dysfunction throughout the basal ganglia and dopamine neuron systems. Furthermore, combining Cre and FlpO knock-in lines will allow for intersectional genetic approaches for simultaneous investigation of neuronal populations. For example, crossing the D1-Cre or A2a-Cre mice with the DAT-FlpO line, could enable dissection of anatomically and/or functionally distinct circuits between the striatum and the VTA and SNc (Lerner et al., 2015; Poulin et al., 2018).

Whereas our knock-in approach minimizes ectopic expression, it is possible that recombinase integration could alter the expression of D1R, A2aR, or DAT at the mRNA or protein level in ways undetectable by our electrophysiology measurements. Furthermore, as low-level expression in cortical and hippocampal regions was observed in D1 and A2a knock-in lines, which is to be expected (Levey et al., 1993; Lammel et al., 2015), it is imperative that future studies take into account the ubiquitous expression of D1Rs, A2aRs, and DATs outside of the basal ganglia. Finally, while our assessments suggest that these lines exhibit largely normal physiological properties, a more comprehensive behavioral panel or additional experimental controls in future experiments are likely necessary to fully confirm that recombinase expression does not alter motor, cognitive, or reward-related behaviors.

In conclusion, these new knock-in lines provide precise genetic tools for interrogating basal ganglia and dopaminergic circuitry: offering improved specificity over BAC transgenic models and support for intersectional genetic strategies. By enabling targeted cell-type manipulations with high specificity, these lines are poised to advance our understanding of basal ganglia function and dysfunction in motor control, reinforcement learning, and neuropsychiatric disorders.

## METHODS

### Data Availability Statement

The data, code, protocols, and key lab materials used and generated in this study are listed in a Key Resource Table alongside their persistent identifiers at 10.5281/zenodo.15499651.

### Contact for reagents and resource sharing

Further information and requests for reagents should be directed to and will be fulfilled by the lead contact, Rui M. Costa.

### Materials availability

The mouse lines generated in this study have been deposited with Jackson Laboratories (D1-Cre – Stock name: *D1-Cre* ; Stock number: 040188. A2a-Cre – Stock name: *A2a-Cre* ; Stock number: 040186. D1-FlpO – Stock name: *D1-FlpO* ; Stock number: 040189. A2a-FlpO – Stock name: *A2a-FlpO* ; Stock number: 040187. DAT-FlpO – Stock name: *DAT-FlpO* ; Stock number: 040184).

### Mouse breeding and husbandry

All experimental protocols were performed according to National Institutes of Health (NIH) guidelines and in compliance with the regulations of the Institutional Animal Care and Use Committee at Columbia University. All experimental animals were 2 to 4 month-old mice housed on a 12 hr light/dark cycle with unrestricted access to food and water.

### Generation of transgenic mice

All knock-in mouse lines were generated by Cyagen (Santa Clara, CA) using CRISPR/Cas9. To generate the D1-Cre mice, a gRNA (5’ GAGATGAGAACCCAATATTC-AGG 3’) targeting exon 2, and a template with a ‘T2A-Cre’ cassette were used. Homology arms were PCR amplified from BAC clones. To generate the A2a-Cre mice, a gRNA (5’ AGGCTGTTCCTACCCTACCC-TGG 3’) targeting exon 3, and a template with a ‘T2A-Cre’ cassette were used. Homology arms were PCR amplified from BAC clones. Two silent mutations, p.A394 (GCC to GCA), and p.R387 (AGG to CGC) were introduced to prevent re-cleavage by Cas9 after homology-directed repair. To generate the D1-FlpO mice, a gRNA (5’ GAGATGAGAACCCAATATTC-AGG 3’) targeting exon 2, and a template with a ‘T2A-FlpO’ cassette were used. To generate the A2a-FlpO mice, a gRNA (5’ AGGCTGTTCCTACCCTACCC-TGG 3’) targeting exon 3, and a template with a ‘T2A-FlpO’ cassette were used. To generate DAT-FlpO mice, a gRNA (5’ CCAACAGCCAATGGCGCAGC-TGG 3’) targeting exon 15, and a template with a ‘2A-FlpO-rBG pA’ cassette were used. Homology arms were PCR amplified from BAC clones RP23-150M11 and RP23-34F24. Two silent mutations, aa 613L (CTG to TTA), and aa 614R (CGC to AGG) were introduced to prevent re-cleavage by Cas9 after homology-directed repair.

### Stereotaxic viral injections

Stereotaxic viral injections Before starting the surgery mice were subcutaneously injected with Buprenorphine XR (0.5–1□mg per kg body weight). The mouse head was shaved, cleaned with 70% alcohol and iodine, an intradermic injection of bupivacaine (2□mg per kg body weight) was administered, and a small incision from anterior to posterior was made on the skin to allow for aligning the head and drilling the hole for the injection site. Surgeries were performed under sterile conditions and isoflurane (1%–5%, plus oxygen at 1-1.5 l/min) anesthesia on a motorized stereotactic frame (David Kopf Instruments, Model 900SD). Throughout each surgery, mouse body temperature was maintained at 37C using an animal temperature controller (ATC2000, World Precision Instruments). To check dopamine neuron expression in the DAT-FlpO mouse, animal were unilaterally injected with 300nL of AAV5-EF1a-fDIO-EYFP (titer: 4.5E12 vg/mL; UNC Vector Core) into the right hemisphere of the substantia nigra pars compacta (SNc; AP -3.16 mm, ML 1.3 mm, DV -4.0 mm) using a Nanoject III Injector (Drummond Scientific, USA) at a pulse rate of 5.1 nL per second, 20 nL per pulse every 5 s. After injection, the pipette was held in place for 5 minutes before raising to ensure minimize viral efflux.

### Slice electrophysiology

2-3 month-old mice were deeply anesthetized with isoflurane (4-5%, oxygen at 1-1.5 l/min) and subsequently transcardially perfused with ice-cold artificial cerebrospinal fluid (ACSF) NMDG cutting solution of the following composition (in mM): 92 NMDG, 2.5 KCl, 1.25 NaH2PO4, 30 NaHCO3, 20 HEPES, 25 glucose, 2 thiourea, 5 Na-ascorbate, 3 Na-pyruvate, 0.5 CaCl2, and 10 MgSO4, pH 7.3-7.4 (Ting et al., 2018). Brains were removed from the skull and glued to the stage of a vibrating microtome (Leica) and 300 μm coronal slices were cut using the same NMDG-ACSF cutting solution. Slices were transferred to a heated chamber at 34 C° with oxygenated NMDG-ACSF recording solution, where they underwent recovery for 25 minutes. Na+ was reintroduced (to 52mM) by gradually adding 2M NaCl-NMDG solution during recovery as previously described (Ting et al., 2018). Slices were then transferred to HEPES-holding solution of the following composition (in mM): 92 NaCl, 2.5 KCl, 1.25 NaH2PO4, 30 NaHCO3, 20 HEPES, 25 glucose, 2 thiourea, 5 Na-ascorbate, 3 Na-pyruvate, 2 CaCl2, and 1 MgCl2, pH 7.3–7.4 at room temperature for at least an hour prior to recordings. Patch electrodes (3–6□MΩ) were filled with a potassium gluconate-based internal solution (135□mM potassium gluconate, 2□mM MgCl2, 0.5□mM EGTA, 2□mM magnesium ATP, 0.5□mM sodium GTP, 10□mM HEPES, 10□mM phosphocreatine and 0.15% Neurobiotin). All recordings were made using a Multiclamp 700B amplifier, the output of which was digitized at 10□kHz (Digidata 1440A). Series resistance was always <35□MΩ and was compensated up to 90%. Neurons were targeted with DIC microscopy and epifluorescence when appropriate. Brain slices were histologically processed to visualize Neurobiotin-filled cells through streptavidin-Alexa Fluor processing. Recording experiments involved presenting a sweep of currents steps (−400pA to 400pA, 20pA step size, 1-second duration). Resulting voltage traces were analyzed using custom MATLAB scripts to extract voltage responses and action potential counts resulting from current injections.

### Histology

Mice were deeply anesthetized with isoflurane and transcardially perfused with PBS, followed by ice-cold 4% paraformaldehyde. Brains were then removed for histological analysis. Coronal or sagittal sections were cut at 75 µm using a Leica VT1000 vibratome. The tissue was rinsed twice in 1x PBS and then permeabilized in PBS containing 0.4% Triton X-100 (PBST). For TH histological labeling, immunohistochemistry was performed with primary antibodies by incubating the sections with TH antibody (Mouse, ImmunoStar, 22941) diluted at 1:2000 in 0.4% Triton X-100 PBS (PBST) overnight at 4C. The sections were then incubated with secondary antibody (Donkey anti-Mouse Alexa Fluor 647, Invitrogen, A31571) diluted at 1:2000 in 0.4% PBST overnight at 4C. DAPI (Sigma D9542; 1:1000) was used as a counterstain in all experiments.

### Image acquisition, processing, and analysis

Coronal or sagittal sections were serially mounted on slides and sections were imaged using an automated slide scanner (Nikon AZ100 Multizoom microscope) equipped with a 4x 0.4NA Plan Apo objective (Nikon Instruments Inc) and P200 slide loader (Prior Scientific), controlled by NIS-Elements using custom acquisition scripts (Nikon Instruments Inc.). Image processing and analysis using BrainJ proceeded as previously described (Botta et al., 2020).

### Statistics

Animal subjects were randomized within experimental blocks to yield equal sampling of experimental conditions. All experiments were conducted in a blind fashion (i.e., experimenter did not know the mouse genotype). Data distribution was assumed to be normal, but this was not formally tested (hence individual data points are presented in all statistical comparisons). Multiple comparisons were conducted with Kruskal-Wallis tests + Dunn’s multiple comparisons test. Unless otherwise specified, two-sided statistical tests were conducted and data is presented as mean ± SEM (standard error of the mean), with all statistical tests, statistical significance values, and sample sizes described in the figure legends.

## Acknowledgments

We would like to thank Zuckerman Institute’s Cellular Imaging platform for providing instrument use and technical advice. Special thanks to Gabriela J. Martins, Mariana L. Correia and Helena M Seuffert for mouse maintenance and Drew Baughman, Mafalda Vicente and Chrissy Weber-Schmidt for additional support with lab and mouse management. This work was supported by grants from the Aligning Science Across Parkinson’s (ASAP-020551) through the Michael J. Fox Foundation for Parkinson’s Research (MJFF), the Parkinson’s Disease Foundation (PDFPF-RCE-1948), the National Institutes of Health (NIH, U19: 5U19NS104649-03), and a NARSAD Young Investigator Grant from the Brain & Behavior Research Foundation (30086). For the purpose of open access, the author has applied a CC BY public copyright license to all Author Accepted Manuscripts arising from this submission.

## Author contributions

E.A. and R.M.C. designed the study, interpreted the results, and wrote the manuscript. A.F., C.C. and L.N. performed histological assessments. E.A. and A.N. conducted electrophysiological characterization. T.S., D.S.P., and R.M.C. supervised the project. All authors contributed to the interpretation of results and manuscript editing.

